# Global Epistasis Makes Adaptation Predictable Despite Sequence-Level Stochasticity

**DOI:** 10.1101/001784

**Authors:** Sergey Kryazhimskiy, Daniel P. Rice, Elizabeth R. Jerison, Michael M. Desai

## Abstract

Epistasis can make adaptation highly unpredictable, rendering evolutionary trajectories contingent on the chance effects of initial mutations. We used experimental evolution in *Saccharomyces cerevisiae* to quantify this effect, finding dramatic differences in adaptability between 64 closely related genotypes. Despite these differences, sequencing of 105 evolved clones showed no significant effect of initial genotype on future sequence-level evolution. Instead, reconstruction experiments revealed a consistent pattern of diminishing returns epistasis. Our results suggest that many beneficial mutations affecting a variety of biological processes are globally coupled: they interact strongly, but only through their combined effect on fitness. Sequence-level adaptation is thus highly stochastic. Nevertheless, fitness evolution is strikingly predictable because differences in adaptability are determined only by global fitness-mediated epistasis, not by the identity of individual mutations.

---

Evolutionary outcomes hinge on the rate and selective effects of new adaptive mutations. Together, these determine a population’s adaptability. This has driven extensive prior work to measure the mutation rate and distribution of fitness effects (DFE) of new mutations (1–8), and to predict how these parameters determine the outcomes of adaptation (9–12). However, recent work in microbial and viral systems shows that epistasis is often pervasive, leading to a rugged fitness landscape where adaptability depends strongly and without any apparent regularity on genotype (13–18). In some cases, just a single mutation can open up previously unavailable opportunities for a population to colonize a new metabolic niche (14) or survive in a previously intolerable drug concentration (15).

This widespread epistasis suggests that the outcomes of adaptation depend strongly on historical contingency (i.e., the specific intermediate genotypes that happen to arise), and are therefore highly unpredictable. On the other hand, others have argued that pervasive but systematic patterns of epistasis can explain striking observations of convergent and parallel evolution at both phenotypic (12, 19, 20) and genotypic levels (21). Related work has provided evidence for such patterns among particular small sets of mutations that arise in a single experimental line (22, 23). These observations suggest a more optimistic picture: evolutionary outcomes may be statistically predictable if mutations leading to extreme and irregular changes in adaptability are rare, while mutations leading to small and regular changes in adaptability are common. Here, we directly test this hypothesis by measuring the variation in adaptability between related genotypes in laboratory yeast populations. We combine this with complete genome sequencing of evolved populations and targeted reconstructions of specific mutations, to connect phenotypic predictability to its genetic causes and to identify the epistatic basis of variation in adaptability.

To this end, we conducted a hierarchical laboratory evolution experiment in *S. cerevisiae* (Figure 1). In the first phase (“Diversification”), we created 432 independent lines from a single haploid clone we refer to as the diversification ancestor (DivAnc). We evolved each line independently in rich media in 96-well microplates for 250 generations, half at large and half at small population size (Methods). We then selected a single clone from 64 of these lines, chosen to span a range of fitness relative to the DivAnc. We refer to these as Founders (Table S1). Each Founder differs by a few mutations, some beneficial and some deleterious. In the second phase (“Adaptation”), we founded 10 independent replicate populations with each Founder, and allowed each of the resulting 640 lines to adapt at a large population size for 500 generations (see Methods). This design allows us to compare variation among lines descended from the same Founder (which reflects the inherent stochasticity of evolution) to variation between lines descended from different Founders, in order to assess the extent to which the genetic background influences future evolution.

**Figure 1.**
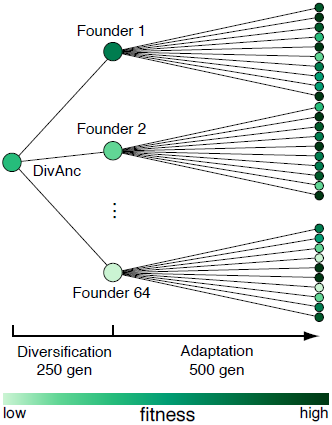
Experimental design. We created many independent lines from a single clone (DivAnc) and evolved each for 250 generations (Diversification). We then selected a single “Founder” clone from 66 of these lines (chosen to span a range of fitness) and evolved 10 independent replicate populations descended from each Founder (Adaptation).

We measured the competitive fitness of each population after 250 and 500 generations of evolution in the Adaptation phase and found that it increased on average by 3.3% and 6.6%, respectively (Figure 2A). However, not all populations adapted at the same rate. Instead, we observe a strong pattern of convergent evolution: the initially large variation in fitness between lines monotonically declined with time (Figure 2A). To quantify the sources of this variation, we first carried out an analysis of variance in the spirit of Travisano *et al*. (24). We partitioned the observed variation in the fitness increase during the Adaptation phase into contributions from measurement noise, inherent stochasticity of the evolutionary process, and the specific genotype of the Founder (Methods). We found that after 250 (500) generations of adaptation, inherent stochasticity explains 49% (29%) of the variance in fitness increment, while 17% (21%) is attributed to measurement error and 34% (50%) to the identity of the Founder (Figure 2B, Table S2). This analysis demonstrates that genetic background is a key determinant of how rapidly a population will adapt. That is, different Founders have dramatically different adaptabilities.

**Figure 2.**
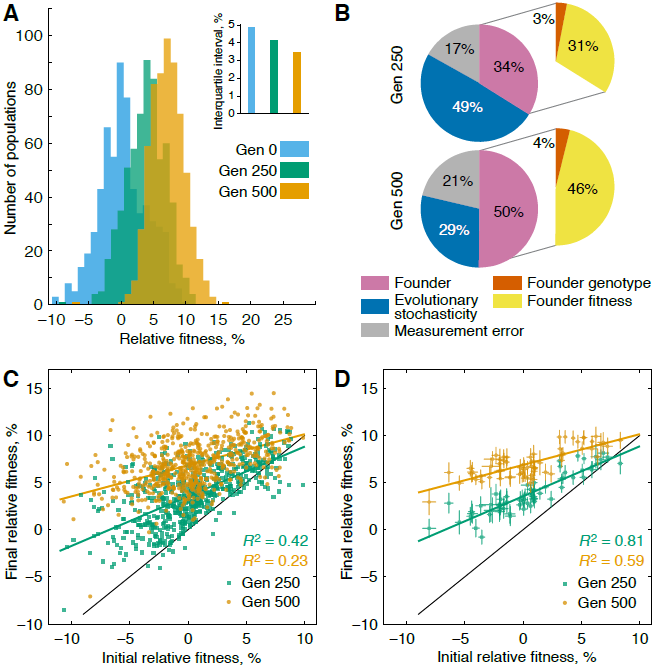
Fitness evolution. **(A)** The distribution of mean population fitnesses over time, relative to DivAnc. Inset shows inter-population fitness variation over time. **(B)** Partitioning of variance in fitness increment after 250 and 500 generations of the Adaptation phase. All variance components are significant (Table S2, Methods). **(C)** Correlation between Founder fitness and population fitness after 250 and 500 generations of Adaptation. **(D)** Correlation between Founder fitness and the mean fitness of the 10 independent lines descended from that Founder, after 250 and 500 generations of Adaptation. Error bars show ±1 sem.

However, these differences in adaptability are not entirely random. Rather, populations with lower initial fitness systematically adapt more rapidly than populations with higher initial fitness, driving the overall pattern of convergent evolution in fitness (Figure 2C). To quantify this effect, we partitioned the variation in fitness increment further: after 250 (500) generations of adaptation, 31% (46%) of this variation is explained by the fitness of the Founder while only 3% (4%) is determined by its specific genotype (Figure 2B, Table S2). In other words, the dramatic differences in adaptability between Founders are almost entirely predicted by their differences in fitness, and independent of the specific mutations that led to that fitness. Thus when we eliminate the effect of inherent stochasticity by averaging the final fitness of populations descended from the same Founder, we find that the initial fitness of the Founder is strongly predictive of the rate of adaptation in its descendant lines (Figure 2D). This means that although inherent evolutionary stochasticity makes it impossible to precisely predict adaptation in individual lines, we can accurately predict the average rate of adaptation based only on the fitness of the initial genotype. We note, however, that although the effects of specific genotype on adaptability are rare or weak, they are significant (Table S2). For example, Founders L041 and L094 have similar fitness but systematic differences in adaptability (Figure S1, Table S3).

A negative correlation between fitness and adaptability has also been observed in prokaryotes among genotypes with fitness variation generated by deleterious mutations (19, 20). It is also consistent with the common observation in many evolution experiments that the rate of increase in fitness within an individual population typically slows down over time (12, 25). Combined with this earlier work, our results suggest a general “rule of declining adaptability” which holds for prokaryotes and eukaryotes adapting to rich and poor media. Further, our observations support an even stronger version of this rule: (a) genotypes with lower fitness are more adaptable than those with higher fitnesses, and (b) distinct genotypes with identical fitness are equally adaptable. We note that this latter observation is inconsistent with Fisher’s geometric model of adaptation (except possibly in one dimension, see Methods and Figure S2).

The rule of declining adaptability could arise for one of two basic (and non-exclusive) reasons. The first explanation is that there are only a few ways to increase fitness, each of which could consist of a single mutation or more generally a set of mutations within some biological module. In this “modular epistasis” model (21), which is a generalization of a finite sites model in population genetics, each beneficial mutation improves a single module and only one mutation is necessary per module. Higher-fitness genotypes adapt more slowly because they have fewer remaining modules to improve.

Alternatively, diminishing returns epistasis may be pervasive among adaptive mutations, such that mutations arising in higher-fitness backgrounds are less beneficial than those arising in lower-fitness backgrounds. This could hold only on average, or it could be true for each mutation individually. Specifically, if epistasis is “idiosyncratic,” mutations may often have widely different effects in different genetic backgrounds (possibly including sign epistasis), but the average effect of a beneficial mutation is smaller in fitter backgrounds. On the other hand, if epistasis is “global,” each individual beneficial mutation provides a smaller advantage in a fitter genetic background. This latter model implies a global coupling among all mutations, such that the effect of each mutation depends on all other mutations, but only through their combined effect on fitness.

To discriminate between the modular, idiosyncratic, and global epistasis models, we sampled one clone from each of 128 populations descended from 15 Founders at generation 500 of the Adaptation phase and sequenced their complete genomes (Methods). Four clones acquired a mutator phenotype during the Adaptation phase and two Founders and all their descendants became diploid (Methods; Figure S3); we excluded these from further analysis, leaving a total of 105 sequenced clones descended from 13 Founders. We identified a total of 55 mutations that occurred in the Founders during Diversification and 1150 mutations that occurred during Adaptation. We annotated each mutation to a gene or intergenic region and classified coding mutations as synonymous or nonsynonymous (Figure 3A, Table S4). Most synonymous and intergenic mutations are likely to be neutral hitchhikers and therefore uninformative with respect to model selection. We therefore restrict further analysis to nonsense, frameshift, nonsynonymous, and promoter mutations, which we refer to as “putatively functional”.

**Figure 3.**
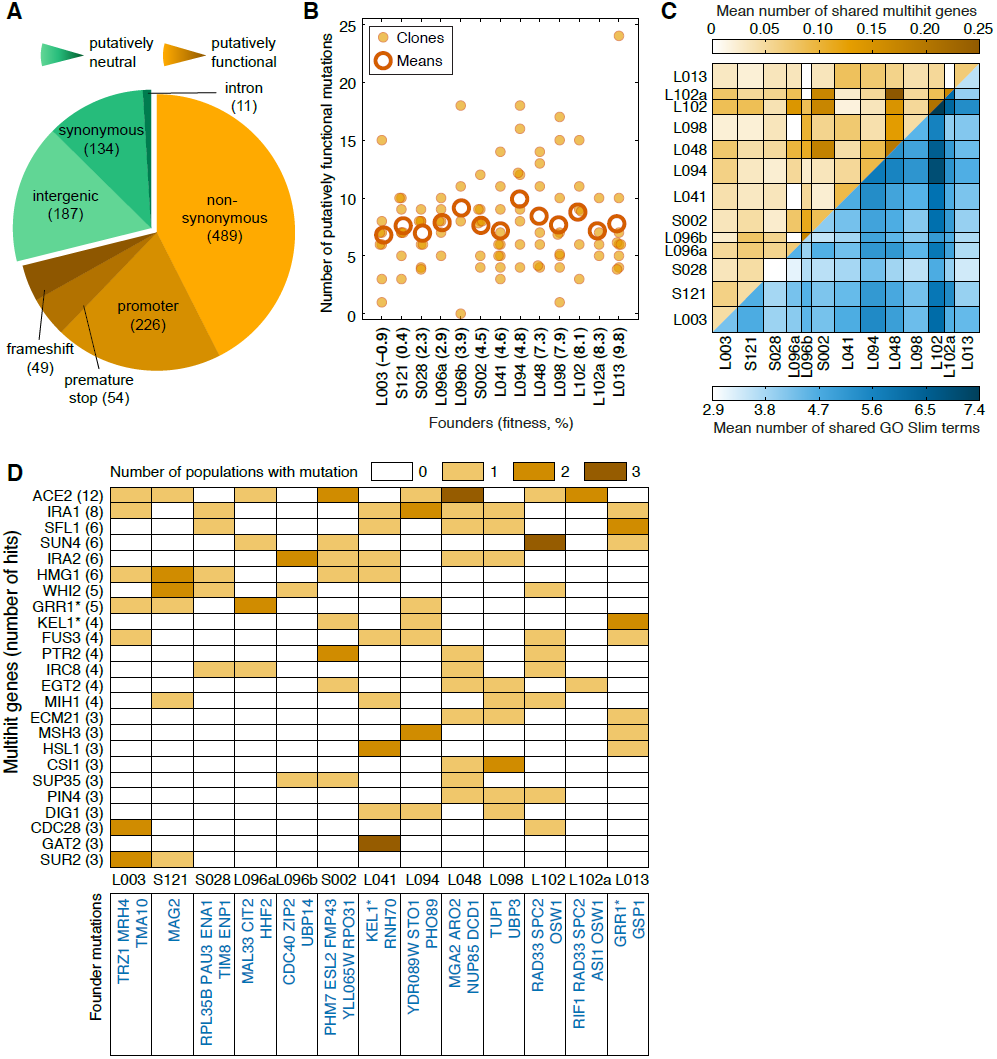
Sequence-level evolution. (**A**) 1150 mutations that occurred in the Adaptation phase arranged by type (Methods). (**B**) Clones descended from different Founders acquired on average the same number of putatively functional mutations (see also Figure S6). (**C**) Convergence and parallelism at the gene (top, orange) and GO Slim (bottom, blue) levels. Each cell is colored to show the average number of mutations shared by two clones descended from the Founders indicated in the row and column headers. Founders are ordered from least-fit (left, bottom) to most-fit (right, top). (**D**) Mutations in multihit genes and the Founder backgrounds in which they were observed (top); putatively functional mutations that determine the Founder background (bottom). Asterisks indicate genes mutated in both Diversification and Adaptation phases.

In contrast to recent experiments in bacteria and viruses (21, 26, 27), all but 4 mutations we identified are unique at the nucleotide level, consistent with our earlier work in *S. cerevisiae* (28). However, we found significant convergent evolution at the gene level. For example, 24 genes had mutations in at least three replicate lines (compared to 4.2 expected by chance, *P* < 10^-4^; Tables S5, S6), indicating that most mutations observed in these multi-hit genes are likely beneficial. Moreover, we observed significant enrichment for mutations in genes involved in negative regulation of Ras, cell cycle regulation, and filamentous growth (Table S7), indicating convergence at higher levels of biological organization. However, it is interesting to note that the degree of convergence in our system is much weaker than recently observed by Tenaillon *et al*. (21) in *E. coli* adapting to heat stress (Figure S4). The biological reasons for this difference are unclear.

We next compared the total number of mutations observed in different evolved lines. As expected, we find that among lines descended from a given Founder, the lines that increased most in fitness were on average those that acquired more mutations in multi-hit genes (Figure S5). In the modular epistasis model, we also expect that lines descended from high-fitness Founders should acquire fewer beneficial mutations than those descended from low-fitness Founders (the former are less adaptable because they have fewer ways to improve). However, we find that this is not the case: lines descended from all Founders acquired the same number of putatively functional mutations (Figure 3B). Similar results hold if we consider only mutations in multi-hit genes (Figure S6).

We next asked whether lines descended from the same Founder took more similar mutational trajectories than lines descended from different Founders. The modular and idiosyncratic epistasis models predict that many mutations are beneficial only in particular genetic backgrounds. Hence, these models predict that clones descended from the same Founder should on average have more mutations in common (“parallelism”) than expected by chance given the observed degree of overall convergence. However, this is not the case. Figures 3C, 3D and S7 show that clones descended from the same Founder are no more likely to share mutations than clones descended from different Founders. This holds regardless of the level at which we define parallelism and convergence (genes or GO Slim categories).

Together, these analyses suggest that the modular and idiosyncratic epistasis models are inconsistent with our data, and point towards the global diminishing returns epistasis model. To confirm this directly, we carried out a series of genetic manipulation experiments. We selected three genes (*SFL1*, *WHI2*, and *GAT2*) in which we found putative loss-of-function (nonsense or frameshift) mutations in three or more replicate lines, suggesting that knockouts of these genes are beneficial in our system. We note that *GAT2* displays the strongest signature of parallel evolution in our data (Figure 3D), and hence represents the strongest candidate for idiosyncratic epistasis. We introduced targeted knockouts of these mutations (along with one control gene, *HO*) separately in several replicates each into all 13 sequenced Founders, as well as in the DivAnc and in four additional clones (Methods, Table S1). We then measured the fitness effects of each knockout in each background. We found a striking negative correlation between the fitness effect of the *gat2*Δ, *whi2*Δ and *sfl1*Δ gene deletions and the fitness of the background strain (Figure 4). Furthermore, there are no idiosyncratic epistatic interactions specific to particular genotypes: up to small deviations, the fitness effect of each knockout depends *only* on the fitness of the genetic background and not on the specific mutations present in that background. This strongly supports the global diminishing returns model as the underlying reason for the rule of declining adaptability with increasing fitness.

**Figure 4.**
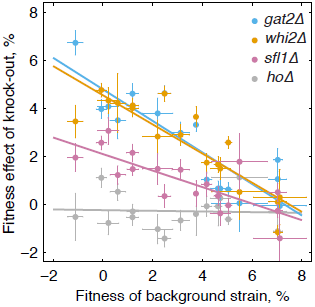
Diminishing returns epistasis among specific mutations. The fitness effect of knocking out genes *gat2*, *whi2*, and *sfl1* declines with the fitness of the background strain. The *ho* knockout is a negative control (Methods). Error bars show ±1 sem over biological replicates.

Taken together, our results suggest a different interpretation of the concept of a “fitness peak” in our system. As is apparent from Figure 2D, all adapting lines appear to be converging in fitness towards a point about 8% to 10% more fit than DivAnc. Consistent with this, all three mutations shown in Figure 4 become approximately neutral in genotypes ∼8% more fit than DivAnc. While this is unlikely to be a true fitness peak, it does represent a fitness level above which adaptation slows substantially. However, this fitness plateau can be reached via many different combinations of beneficial mutations. That is, it occupies not a single point in genotype space but rather an enormous number of genetically and perhaps even physiologically distinct types. A population adapting on this fitness landscape may take one of many thousands of genetically distinct trajectories, each of which would nevertheless lead it to the same fitness plateau at the same rate.

These results paint a surprisingly simple picture of adaptation in our system. Many mutations scattered across many biological processes appear to be beneficial. Yet despite their lack of apparent functional relationship, these mutations are globally coupled by diminishing returns epistasis – their effects are strongly mediated by background fitness, but are otherwise essentially independent of the specific identity of other mutations present in the background. The biological basis of this global coupling remains unknown. Nevertheless, it leads to a striking pattern of convergent evolution at the fitness level, making fitness evolution relatively predictable. Despite this fitness-level convergence, evolution remains highly stochastic at the genotype level, because many different mutations can be responsible for a given fitness change.

## Acknowledgements

We thank Andrew Murray, Melanie Müller, Benjamin Good, David van Dyken, Michael McDonald, and other members of the Desai lab for useful discussions and comments on the manuscript; Arvind Subramaniam, Greg Lang, Melanie Müller, and John Koschwanez for experimental advice and strains; and Patricia Rogers and Christian Daly for technical support. This work was supported by a Career Award at the Scientific Interface from the Burroughs Wellcome Foundation (S.K.), two NSF graduate research fellowships (D.P.R., E.R.J.), and by the James S. McDonnell Foundation, the Alfred P. Sloan Foundation, the Harvard Milton Fund, grant PHY 1313638 from the NSF, and grant GM104239 from the NIH (M.M.D.). Computational work was performed on the Odyssey cluster supported by the Research Computing Group at Harvard University.

## References Cited

1. S. B. Joseph, D. W. Hall, Spontaneous Mutations in Diploid Saccharomyces cerevisiae: More Beneficial Than Expected. Genetics 168, 1817 (December 1, 2004, 2004).

2. L. Perfeito, L. Fernandes, C. Mota, I. Gordo, Adaptive mutations in bacteria: High rate and small effects. Science 317, 813 (2007).

3. R. C. MacLean, A. Buckling, The Distribution of Fitness Effects of Beneficial Mutations in *Pseudomonas aeruginosa*. PLoS genetics 5, e1000406 (2009).

4. M. Imhof, C. Schlotterer, Fitness effects of advantageous mutations in evolving Escherichia coli populations. Proc. Natl. Acad. Sci. USA 98, 1113 (2001).

5. R. Kassen, T. Bataillon, Distribution of Fitness Effects Among Beneficial Mutations Before Selection in Experimental Populations of Bacteria. Nature Genetics 38, 484 (2006).

6. R. Sanjuan, A. Moya, S. Elena, The distribution of fitness effects caused by single-nucleotide substitutions in an RNA virus. Proc. Natl. Acad. Sci. USA 101, 8396 (2004).

7. D. R. Rokyta, An empirical test of the mutational landscape model of adaptation using a single-stranded DNA virus. Nat. Genet. 37, 441 (2005).

8. D. E. Rozen, J. A. de Visser, P. J. Gerrish, Fitness effects of fixed beneficial mutations in microbial populations. Curr. Biol. 12, 1040 (2002).

9. P. Gerrish, R. Lenski, The Fate of Competing Beneficial Mutations in an Asexual Population. Genetica 102/103, 127 (1998, 1998).

10. P. D. Sniegowski, P. J. Gerrish, Beneficial mutations and the dynamics of adaptation in asexual populations. Philosophical Transactions of the Royal Society B: Biological Sciences 365, 1255 (April 27, 2010, 2010).

11. B. H. Good, I. M. Rouzine, D. J. Balick, O. Hallatschek, M. M. Desai, The rate of adaptation and the distribution of fixed beneficial mutations in asexual populations. Proceedings of the National Academy of Sciences 109, 4950 (2012).

12. M. J. Wiser, N. Ribeck, R. E. Lenski, Long-Term Dynamics of Adaptation in Asexual Populations. Science, (November 14, 2013, 2013).

13. R. J. Woods et al., Second-Order Selection for Evolvability in a Large Escherichia coli Population. Science 331, 1433 (March 18, 2011, 2011).

14. Z. D. Blount, C. Z. Borland, R. E. Lenski, Historical Contingency and the Evolution of a Key innovation in an Experimental Population of E*scherichia coli*. PNAS 105, 7899 (2008).

15. J. D. Bloom, L. I. Gong, D. Baltimore, Permissive Secondary Mutations Enable the Evolution of Influenza Oseltamivir Resistance. Science 328, 1272 (June 4, 2010, 2010).

16. D. R. Rokyta et al., Epistasis between Beneficial Mutations and the Phenotype-to-Fitness Map for a ssDNA Virus. PLoS genetics 7, e1002075 (2011).

17. C. L. Burch, L. Chao, Evolvability of an RNA virus is determined by its mutational neighbourhood. Nature 406, 625 (2000).

18. D. M. Weinreich, N. F. Delaney, M. A. DePristo, D. L. Hartl, Darwinian Evolution Can Follow Only Very Few Mutational Paths to Fitter Proteins. Science 312, 111 (2006).

19. J. E. Barrick, M. R. Kauth, C. C. Strelioff, R. E. Lenski, Escherichia coli rpoB Mutants Have Increased Evolvability in Proportion to Their Fitness Defects. Molecular Biology and Evolution 27, 1338 (June 1, 2010, 2010).

20. L. Perfeito, A. Sousa, T. Bataillon, I. Gordo, Rates of Fitness Decline and Rebound Suggest Pervasive Epistasis. Evolution, n/a (2013).

21. O. Tenaillon et al., The Molecular Diversity of Adaptive Convergence. Science 335, 457 (January 27, 2012, 2012).

22. A. I. Khan, D. M. Dinh, D. Schneider, R. E. Lenski, T. F. Cooper, Negative Epistasis Between Beneficial Mutations in an Evolving Bacterial Population. Science 332, 1193 (June 3, 2011, 2011).

23. H.-H. Chou, H.-C. Chiu, N. F. Delaney, D. Segrè, C. J. Marx, Diminishing Returns Epistasis Among Beneficial Mutations Decelerates Adaptation. Science 332, 1190 (June 3, 2011, 2011).

24. M. Travisano, J. Mongold, A. Bennett, R. Lenski, Experimental tests of the roles of adaptation, chance, and history in evolution. Science 267, 87 (January 6, 1995, 1995).

25. S. Kryazhimskiy, G. Tkačik, J. B. Plotkin, The dynamics of adaptation on correlated fitness landscapes. Proceedings of the National Academy of Sciences 106, 18638 (November 3, 2009, 2009).

26. J. P. Bollback, J. P. Huelsenbeck, Clonal Interference Is Alleviated by High Mutation Rates in Large Populations. Mol Biol Evol 24, 1397 (June 1, 2007, 2007).

27. A. J. Betancourt, Genomewide Patterns of Substitution in Adaptively Evolving Populations of the RNA Bacteriophage MS2. Genetics 181, 1535 (April 1, 2009, 2009).

28. G. I. Lang et al., Pervasive genetic hitchhiking and clonal interference in forty evolving yeast populations. Nature 500, 571 (2013).

